# The 3D Genome Browser: a web-based browser for visualizing 3D genome organization and long-range chromatin interactions

**DOI:** 10.1101/112268

**Authors:** Yanli Wang, Bo Zhang, Lijun Zhang, Lin An, Jie Xu, Daofeng Li, Mayank NK Choudhary, Yun Li, Ming Hu, Ross Hardison, Ting Wang, Feng Yue

## Abstract

Recent advent of 3C-based technologies such as Hi-C and ChIA-PET provides us an opportunity to explore chromatin interactions and 3D genome organization in an unprecedented scale and resolution. However, it remains a challenge to visualize chromatin interaction data due to its size and complexity. Here, we introduce the 3D Genome Browser (http://3dgenome.org), which allows users to conveniently explore both publicly available and their own chromatin interaction data. Users can also seamlessly integrate other “omics” data sets, such as ChIP-Seq and RNA-Seq for the same genomic region, to gain a complete view of both regulatory landscape and 3D genome structure for any given gene. Finally, our browser provides multiple methods to link distal *cis*-regulatory elements with their potential target genes, including virtual 4C, ChIA-PET, Capture Hi-C and cross-cell-type correlation of proximal and distal DNA hypersensitive sites, and therefore represents a valuable resource for the study of gene regulation in mammalian genomes.

---

The three-dimensional (3D) organization of mammalian genomes has been shown to be essential in gene regulation^1-4^. At the DNA level, distal regulatory elements such as enhancers need to be in physical contact with their target genes. At a larger scale, topologically associating domains (TADs) have been suggested to be the basic unit of mammalian genome organization^5^. Several recent high-throughput technologies based on Chromatin Conformation Capture (3C)^6^ have emerged (such as Hi-C^7^, ChIA-PET^8^, Capture-C^9^, Capture Hi-C^10^, PLAC-Seq^11^ and HiChIP^12^) and have provided an unprecedented opportunity to study this spatial organization in a genomewide fashion.

As the volume of chromatin interaction data generated from individual labs and large collaborative projects such as the ENCODE^13^ and 4D Nucleome consortia increases, efficient visualization and navigation of these data becomes a major bottleneck for their biological interpretation. Due to the size and complexity of these interactome data, it is difficult to store and explore them on a personal device. In tackling this challenge, several useful tools for visualizing chromatin interactions have been developed, and each of them has its unique features and limitations. The Hi-C Data Browser is the first web-based query tool that visualizes Hi-C data as heatmaps. Unfortunately, it does not support zoom functionalities and only hosts limited number of datasets. The WashU Epigenome Browser^14,15^ can display both Hi-C and ChIA-PET data, and it provides access to thousands of epigenomic datasets from the ENCODE and Roadmap Epigenome projects. Although the WashU Epigenome Browser allows users to upload and visualize their own datasets, it may be challenging to take advantage of this feature as the file size of Hi-C matrices, especially high-resolution ones, could reach hundreds of gigabytes. Juicebox^16^, a Java program that was introduced as a companion tool for exploring the high-resolution Hi-C data made available by the initial paper^17^, has many practical features, such as zoom functionalities and supplementation of other epigenomic data. It is, however, not portable, as it is not web-based and requires installation. In addition, most aforementioned tools only display Hi-C as a heatmap, which is convenient for visualizing large domain structures such as TADs, but may not be the most intuitive way for visualizing enhancer-promoter interactions.

Here, we present the 3D Genome Browser (www.3dgenome.org), which is a fast and portable web-based browser that allows users to smoothly explore both published and their own chromatin interaction data. Our 3D Genome Browser features five display modes: 1) Hi-C contact matrices as heatmaps along with corresponding TAD information, 2) stacked Hi-C heatmaps to facilitate cross-tissue or even cross-species comparisons of chromatin structures between two different Hi-C datasets, 3) Hi-C-derived virtual 4C plot along with ChIA-PET data, 4) ChIA-PET or other chip-based chromatin interaction data such as PLAC-Seq and HiChIP, and 5) Capture Hi-C or other capture-based chromatin interaction data. These distinct and versatile display modes encourage users to explore interactome data tailored to their own needs, from exploring organization of higher-order chromatin structures to investigating enhancer-promoter interactions. Our browser provides zoom and traverse functionalities in real time and supports querying inputs of genomic features such as gene name or SNP rsid. The 3D Genome Browser provides users with the ability to highlight loci of interest and to zoom into or center onto these regions as needed. In addition, the browser incorporates the UCSC Genome Browser and the WashU Epigenome Browser, allowing users to simultaneously query and supplement chromatin interaction data with thousands of genetic, epigenetic and phenotypic datasets, including ChIP-Seq and RNA-Seq data from the ENCODE and Roadmap Epigenomics projects. The overall structure of the 3D Genome Browser is summarized in **Fig. 1**.

**Figure 1.**
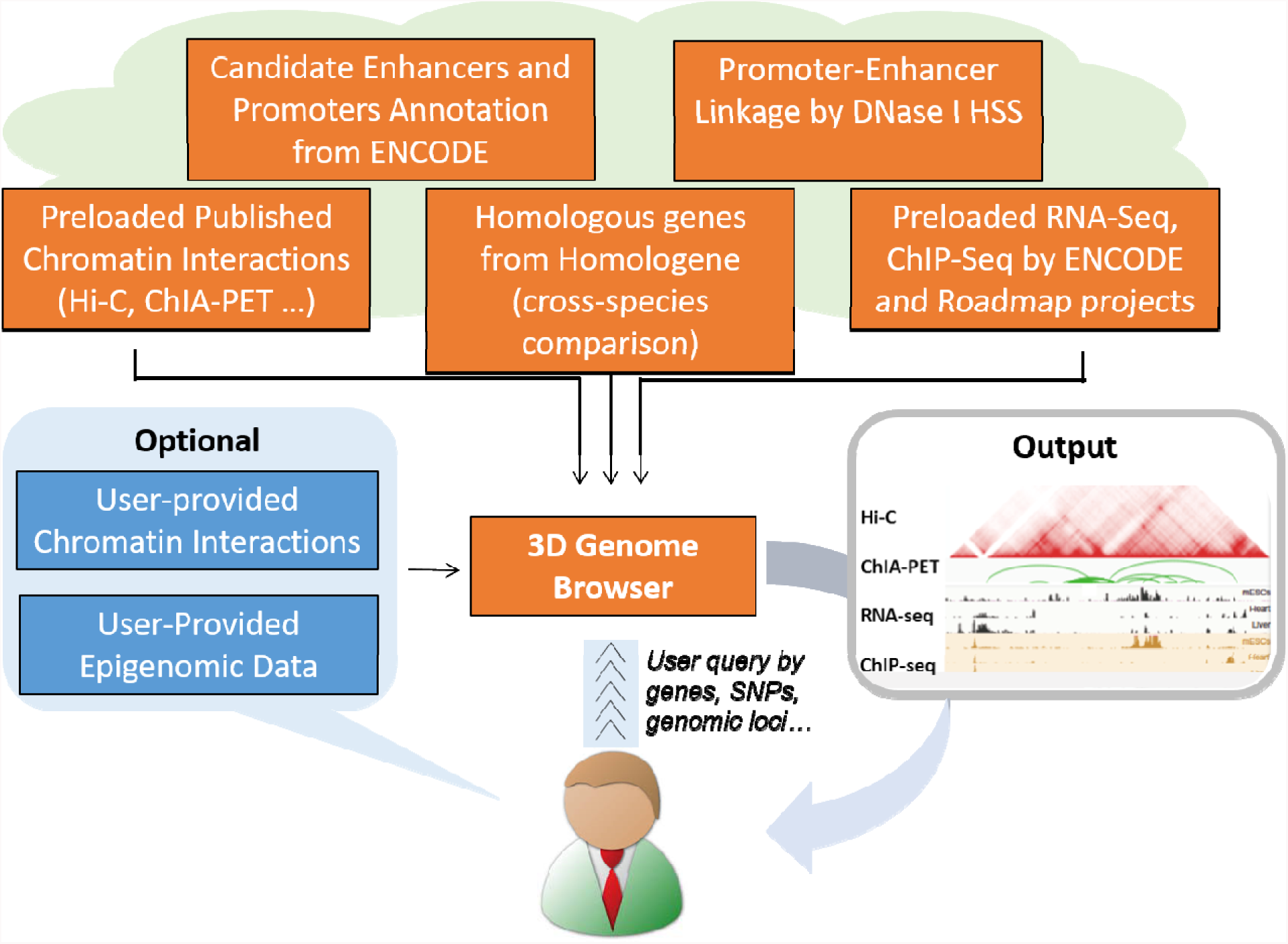
Overall design of the 3D Genome Browser.

## Hi-C Heatmap

In this mode, Hi-C contact matrices are visualized as heatmaps aligned with an embedded external browsers (UCSC or WashU). Our users may choose to display 86 Hi-C datasets of different resolution (matrix bin-sizes) from multiple human and mouse tissues, including 12 that were recently remapped to more updated genome assembly (hg18→hg19). In addition, we predicted TADs for all datasets using streamlined pipeline based on Dixon *et al*^5^, so that users can conveniently explore domain structures estimated by Hi-C data. Upon data selection, the user can query any region of interest by submitting either genomic coordinates or specific genomic features, such as gene symbol, RefSeq ID, Ensembl ID, Uniprot ID and SNP rsid. As an example, we queried the chromatin interactions for the *SLC25A37* gene (**Fig. 2**) using the 5-kb resolution Hi-C contact map from human K562 cell line^17^. Once the heatmap loads, we could zoom into the region by clicking on the *Zoom In* button or double-clicking on the heatmap. Furthermore, we could adjust the minimum and maximum cutoff values using the slider handles on the color bar to increase the coloring contrast. We observed a heatmap cell with high contact value. To interpret the biological meaning of the Hi-C heatmap, we integrated the WashU Epigenome Browser with gene annotation and histone modification H3K4me1, H3K4me3 and H3K27ac for K562. By single-clicking on the heatmap cell of interest, we determined that the two interacting loci are the promoter of *SLC25A37* and a putative enhancer as determined by histone modification patterns and chromHMM and that the enhancer-promoter pair resides in the same TAD (**Fig. 2**). This putative enhancer has been confirmed to exhibit enhancer activities that regulate *SLC25A37* expression during late phase erythropoiesis^18^.

**Figure 2.**
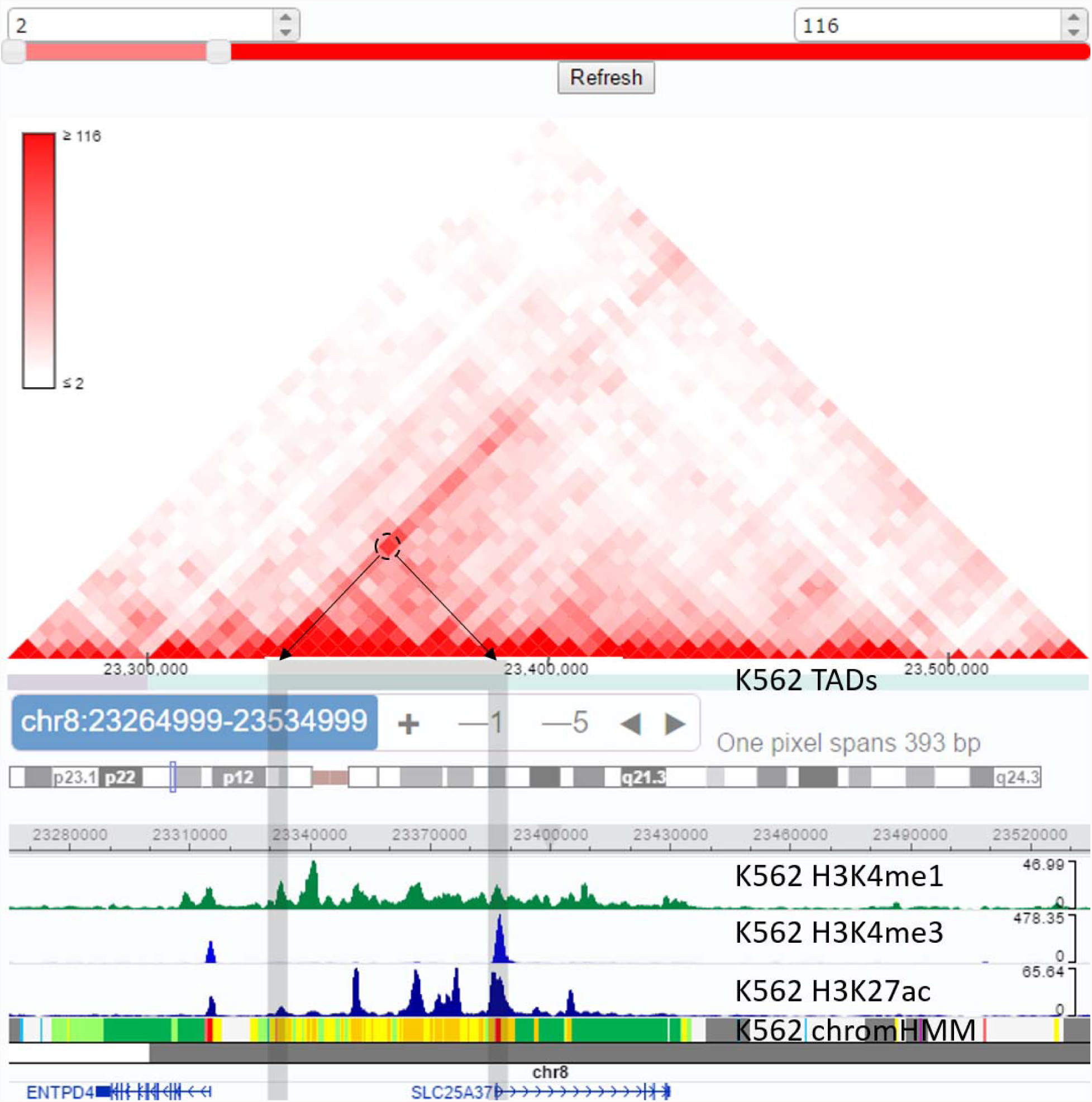
Using the Hi-C mode of the 3D Genome Browser to view chromatin interactions. Browsing the region around the *SLC25A37* in human K562 at 5-kb resolution to find chromatin interactions. Here, we employ the WashU Epigenome Browser as our embedded external browser and obtain gene annotation, ChIP-Seq of H3K4me1, H3K4me3 and H3K27ac histone modifications for K562 as well as its chromHMM data for the corresponding region. We click on a heatmap cell with high value to highlight the two interacting loci, revealing physical contacts between the promoter of *SLC25A37* and a putative enhancer upstream, whose functional linkage was confirmed^18^.

Hi-C heatmaps not only describe chromatin interactions, but also may denote structural variations. Certain structural variations, such as deletion, insertion, inversion and translocation, establish signature patterns within the heatmaps and therefore could be discovered with Hi-C. In our example, we examine a deletion specific to human K562 which involves the genes *CDKN2A* and *CDKN2B* (**Fig. 3a**), as previously characterized^19^. Evidently, our browser could aid in the investigation of structural variations and their impact on chromatin organization in cancers and other diseases.

**Figure 3.**
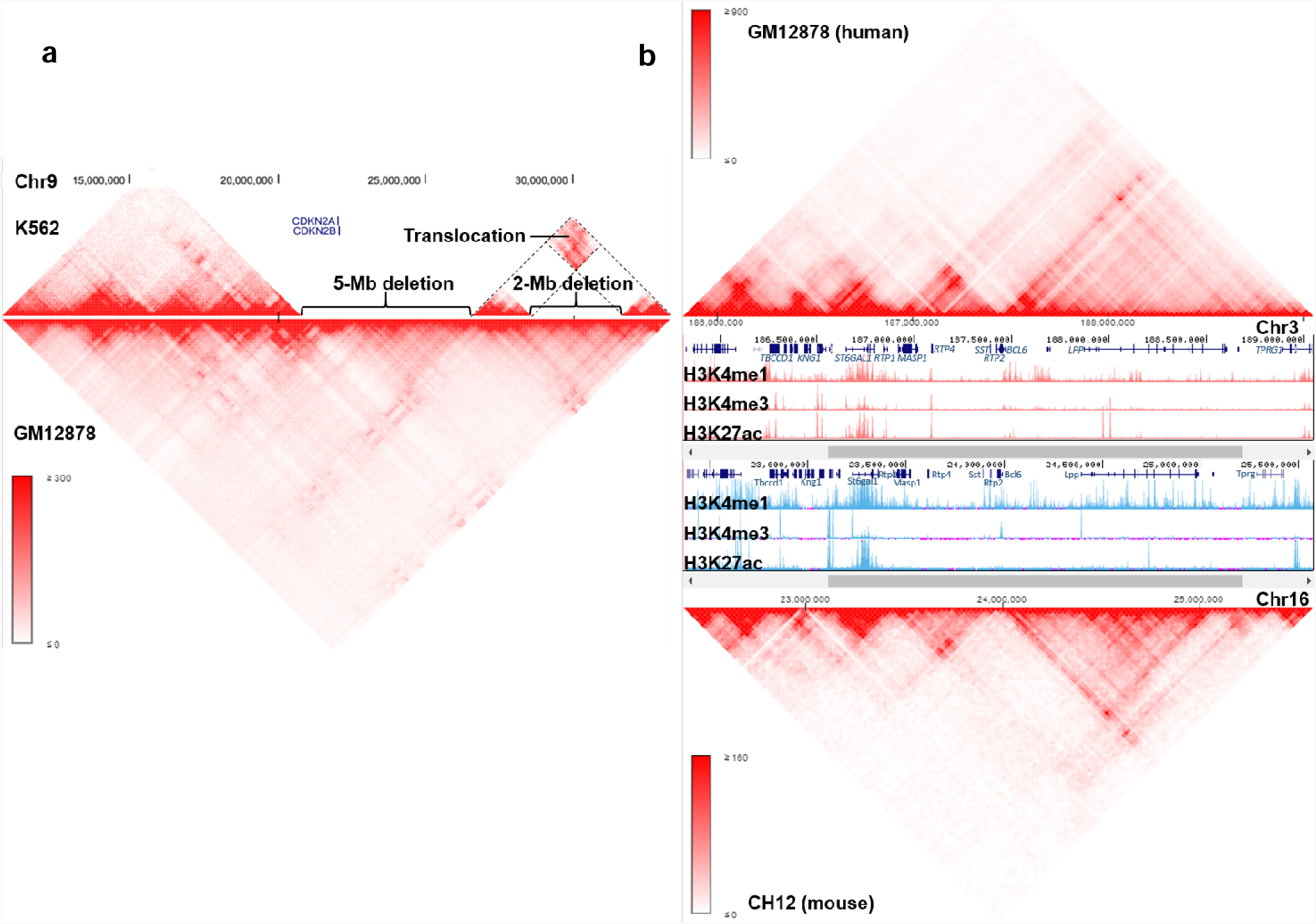
Using the compare Hi-C mode of the 3D Genome Browser for cross-tissue and crossspecies comparisons of Hi-C data: a. cross-tissue comparison of the same species. Comparing human K562 to human GM12878 yields a deletion encompassing *CDKN2A* and *CDKN2B* specific to K562 on chromosome 9; **b. cross-species comparison.** Comparing human GM12878 to mouse CH12 at the region surrounding the *BCL6*/Bcl6 region demonstrates an evolutionary conservation of the chromatin structure between the two species.

## Compare Hi-C

This mode compares two Hi-C datasets by displaying them as two opposing heatmaps stacked along aligned genomic coordinates. This arrangement allows the users to contrast the similarities and differences of chromatin structure between different cells/tissues or different species. When displayed on the same genome assembly, the browser accepts the same inputs as regular Hi-C mode and users can visually distinguish cell-specific chromatin interactions and structural variations (**Fig. 3a**). In contrast, when displayed on assemblies of two different species, the users enter a gene, and the browser will obtain homologous counterpart from the other selected species based on NCBI’s HomoloGene database^20^. This across-species comparison tool would be invaluable in investigating the evolutionary conservation of chromatin organization (**Fig. 3b**).

## Virtual 4C

Although displaying Hi-C data as a heatmap is informative to visualize large genome structures such as TADs, it is not an intuitive way to highlight contacts between two loci of interest, such as enhancer-promoter interactions. To attain loci-specific interactions, we implemented Hi-C-derived virtual 4C plot in our browser. The 4C (Circular Chromosomal Conformation Capture^21,22^) experiment is a chromatin ligation-based method that surveys for *one-vs-many* interactions in the genome, that is, to measure the interaction frequencies between a “bait” locus and any other loci in its vicinity. Its data is plotted as a line histogram, where the center is the “bait” region and any peak signals in distal regions indicate frequency of chromatin interaction events. In our browser, we use the queried region as bait (gene name, SNP ID or genomic coordinates), and extract Hi-C data centered on the bait region, hence, *virtual* 4C.

In addition to the virtual 4C plot, our browser also displays ChIA-PET and DNase I hypersensitive site (DHS)-linkage data to help users hypothesize potential connection between enhancers and their target genes. ChIA-PET is another implementation of chromatin ligation-based method, which detects long-range interactions between genomic regions that are enriched for a feature (either histone modification or transcription factor binding). In our browser, each pairwise ChIA-PET interaction is visualized as an elliptical arc. DHS-linkage is another method linking distal regulatory element with their target genes by computing Pearson correlation coefficients between the gene proximal and distal DHS pairs across more than 100 ENCODE cell types^23^. Similar to ChIA-PET, each associated proximal-distal DHS pair is virtualized as an elliptical arc, with all arcs involving the same proximal DHS being in the same color. Both ChIA-PET and DHS-linkage plots supplement the virtual 4C plot, providing additional layers of evidence for potential enhancer-promoter interaction.

To illustrate the utility of this browser mode, we show an example by querying rs12740374 (**Fig. 4**). The SNP that has been associated with high plasma low-density lipoprotein cholesterol (LDL-C)^24^, which could lead to coronary artery disease and myocardial infarction. Since LDLs are processed by the liver, we examine the histone modifications in the Hep2G cell line and we use virtual 4C and ChIA-PET data from K562, since high resolution Hi-C and numerous ChIA-PET data are only available for K562, but not for hepatic cell lines. The rs12740374 SNP is located within a candidate enhancer region as marked by H3K27ac shown in the UCSC genome browser. In this case, virtual 4C, ChIA-PET and DHS-Linkage all support that there is a putative interaction between the enhancer harboring this SNP and the promoter region of *SORT1*, which has been previously discovered^25^. By integrating multiple lines of evidence, our browser provides a valuable resource for investigators to create hypotheses connecting distal non-coding variant, distal regulatory element and their target gene.

**Figure 4.**
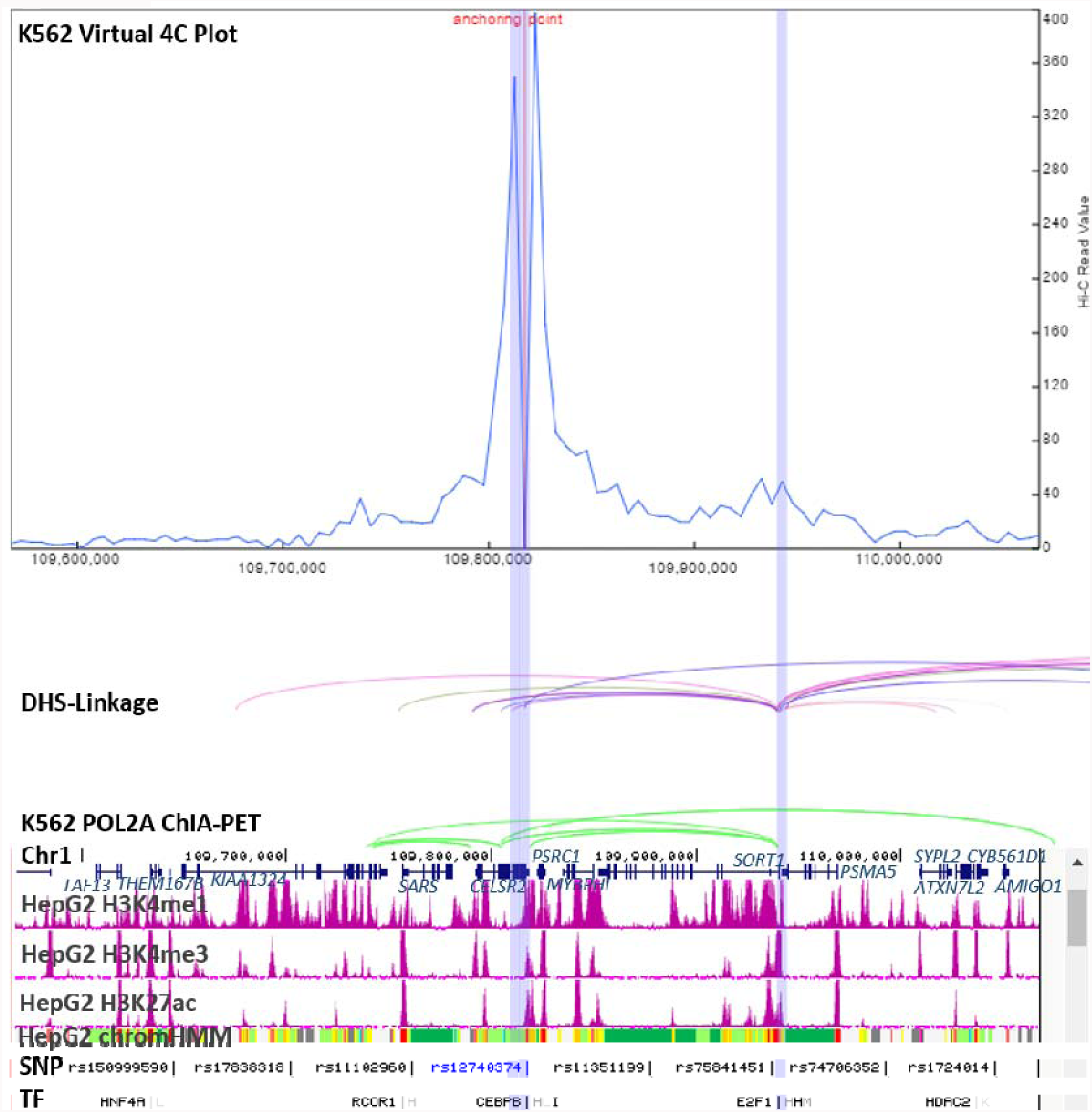
Using the virtual 4C mode of the 3D Genome Browser to identify interactions between cis-regulatory elements (enhancers) and their target genes. Here, we investigate the effect of SNP rs12740374 by plotting the virtual 4C from the 5-kb K562 Hi-C, DHS-linkage and K562 POL2A ChIA-PET and integrating the UCSC Genome Browser. Based on the HepG2 chromHMM, the SNP resides at a putative enhancer region (orange) while according to virtual 4C (not available for HepG2, but the K562 dataset was utilized based on the stability of chromatin structure across cell lines), there is a peak indicating an interaction between this enhancer and the promoter of *SORT1* (transcribed right to left). This is supported by K562 POL2A ChIA-PET (again not available for HepG2) and DHS-linkage data. Based on transcription factor ChIP data, the candidate enhancer has a CEBPB binding site in HepG2 (L), which is possibly disrupted by rs12740374, ultimately affecting *SORT1* expression. All of the above has been confirmed by Musunuru *et al*^24^.

## ChIA-PET and other chip-based methodologies

Along with ChIA-PET, data from other chip-based technologies, such as HiChIP and PLAC-Seq are displayed together with DHS-linkage in this browser mode. Each interaction pair is rendered as an elliptical arc and once queried through a gene, SNP or genome coordinate, could be explored by users alongside their external genome browser (UCSC or WashU) of choice.

## Capture Hi-C and other capture-based techniques

Capture-based chromatin ligation-based methods seek long-range interaction which involves selected elements of interests captured with pre-determined sequences, such as promoters. In this browser mode, Capture Hi-C interactions are visualized similar to ChIA-PET, as elliptical arcs. In our example, we queried the gene *PAX-5* (**Fig. 5**), which is instrumental in B-cell development, in the naïve B-cell dataset^26^. We notice several interactions between the *PAX5* promoter and H3K27ac-rich region downstream of the *ZCCHC7* gene. One region marked by histone modification was indeed determined to be an enhancer for *PAX5* and its disruption leads to leukemogenesis^27^. Once again, the 3D Genome Browser demonstrates its power for hypothesis generation about the potential *cis*-regulatory elements and their potential target genes.

**Figure 5.**
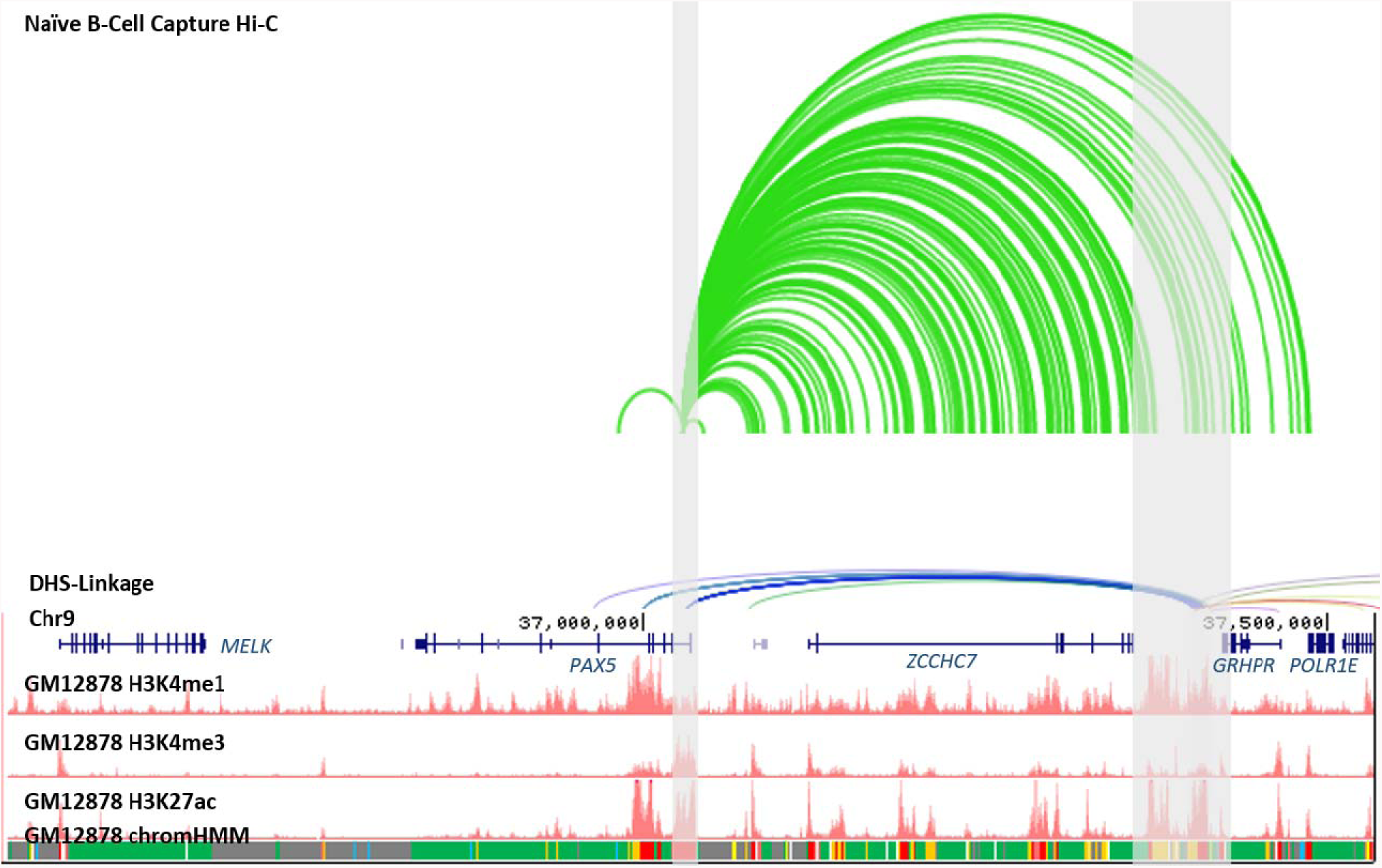
Using the Capture Hi-C mode of the 3D Genome Browser to further dissect enhancer-promoter interactions. We examine the region surrounding *PAX5* in naïve B cells with Capture Hi-C plot and notice a region downstream of *ZCCHC7* rich in H3K4me1 and H3K27ac and classified as putative enhancer by chromHMM (GM12878). Both the Capture Hi-C data and DHS-linkage support an interaction between this possible enhancer and the promoter of *PAX5* (transcribed right to left), which has been confirmed^26^.

## Use Your Own Data and New Data Format

The 3D Browser supports a variety of features that allow users to browse their own or others’ unpublished data. First, our browser encourages integration with customized UCSC sessions, in which the users could add or modify existing tracks or upload their own data to UCSC. To view a customized UCSC session, the user would only be required to enter the session URL. Next, the users could view their own Hi-C data by converting the *n* × *n* contact matrices, where *n* is the number of interrogated loci, into a novel, indexed binary file format called <u>B</u>inary <u>U</u>pper <u>T</u>riangu<u>L</u>ar <u>M</u>at<u>R</u>ix (BUTLR file) developed by us. By hosting the BUTLR file on any HTTP-supported server and providing the URL to the 3D Genome Browser, a user can take full advantage of the features of our browser, without uploading their Hi-C data since the browser would only query the selected region through binary indexing, rather than searching through the entire matrix. This capability is similar to the bigWig/bigBed mechanism invented by UCSC^28^.

Additionally, BUTLR format dramatically reduces the storage of high-resolution Hi-C data not only through the binarization but also through the omission of redundant values (**Supplemental Fig. 1**). Most notably, while 1-kb resolution hg19 intrachromosomal Hi-C contact matrices in the tab-delimited format requires almost 1 TB, the BUTLR format of those same matrices would only need 11GB. The binary file format also greatly improves the query speed: using pre-loaded Hi-C data sets, our 3D browser generally return the query results as a heatmap in the matter of seconds.

In summary, we developed an interactive 3D genomes browser that is being used by thousands of users each month. By simultaneously displaying the 3D chromatin interactions, functional genomic annotations, and disease/trait-associated SNPs, we provide an invaluable online tool to investigators from all over the world for the study of 3D genome organization and its functional implications on gene regulation in mammalian genomes.

## ACKNOWLEDGEMENTS

This work was supported by NIH grant 1U01CA200060. FY is also supported by grants from Leukemia Research Foundation, PhRMA Foundation, and Penn State CTSI. TW is also supported by NIH grants R01HG007175, R01HG007354, R01ES024992, U24ES026699, and U01HG009391. MH is partially supported by NIH U54DK107977 (contact PI: Dr. Bing Ren). YL is partially supported by NIH R01HG006292 and R01HL129132.

